# Impact of Temporal Uncertainty on Sign-tracking Behavior

**DOI:** 10.1101/2024.11.30.626192

**Authors:** Rie Kaneko, Eleanor H Simpson, Peter D Balsam

**Affiliations:** Columbia University

**Keywords:** "Sign-tracking", "Reinforcement learning", "Pavlovian conditioning", "Uncertainty", “Autoshaping”

## Abstract

Sign-tracking behavior, also known as “autoshaping”, is defined as approach and interaction with reward-predictive cues. It is associated with addiction-related phenotypes and compulsive behavior. Several previous studies have demonstrated that when there is uncertainty about reward properties (e.g. probability, size), sign-tracking is increased. However, the effect of cue-uncertainty on sign-tracking behavior is not known. Here, using a Pavlovian conditioning paradigm, we manipulated the temporal uncertainty about the appearance of cues by implementing either fixed or variable inter-trial intervals (ITIs) of different durations across groups of mice. We found that temporal uncertainty during acquisition significantly enhances sign-tracking, which persists during extinction, even when ITI variability was different in the extinction session than in the acquisition session. This suggests that the effects of temporal uncertainty are learned and retained, rather than performance-based. Our results demonstrate that sign-tracking behavior is not only modified by the characteristic of the reward, but it can also be modified by the uncertainty regarding cues. These findings highlight how temporal predictability shapes cue-directed behaviors and has implications for understanding reward-related behavioral responses including sign-tracking behaviors.

## Introduction

Associative learning is a fundamental biological process that enables animals to predict and react to potential outcomes based on environmental cues (Enquist et al., 2016). Crucial for survival, this type of learning promotes adaptive behaviors in response to signals of impending rewards or threats. When the anticipatory response to these cues includes attraction, approach and interaction with the cues (conditioned stimuli, CS) that predict outcomes, the response is called “Sign-tracking”. Attraction, approach and interaction with the location at which outcomes (unconditioned stimuli, US) are presented is described as “Goal-Tracking” (Boakes, 1977; Flagel et al., 2009). These two distinct behaviors develop over the course of associative learning and individual differences in how these behaviors develop may be important for understanding both healthy and pathological reward related behaviors including addictions (Colaizzi et al., 2020).

Animals that display increased sign tracking behavior have a greater addiction vulnerability. For example, sign-trackers work harder to receive cocaine injections (Saunders & Robinson, 2011), exhibit more pronounced cocaine-induced reinstatement compared to goal-trackers even after extinction (Saunders & Robinson, 2010), and show greater propensity for psychomotor sensitization to stimulants F(Flagel et al., 2009). In humans, a greater sensitivity to cues associated with drugs has been documented among regular drug users, such as those who use alcohol (Cofresí et al., 2019; Grüsser et al., 2004; Schacht et al., 2013), cocaine (Kosten et al., 2006; Negrete & Emil, 1992; Robbins et al., 1999), and opioids (Back et al., 2014). Thus, it is important to understand the factors that give these cues the power to strongly evoke attention, approach, and contact.

One factor that increases the salience of these cues is the uncertainty about the predictability of rewards following the cues. Studies indicate that uncertainty regarding whether a CS will be followed by a reward leads to an increase in sign-tracking behavior (Boakes, 1977; Davey & Cleland, 1982; Gibbon et al., 1980) and is maximum when uncertainty is greatest, which is when the probability of being rewarded is 50%, compared to 25% or 75% reward probability (Anselme et al., 2013; Monosov & Hikosaka, 2013; Robinson et al., 2019). Additionally, uncertainty about reward magnitude has been shown to enhance sign-tracking (Anselme et al., 2013; Robinson et al., 2014).

Given that sign-tracking is thought to result from an increase in attention and focus on reward predictive cues, it seems paradoxical that the more uncertain the reward cue is as a source of information about impending reward values, the greater the focus and attraction to that cue. One possible explanation for uncertainty increasing sign-tracking is an evolutionary adaptation to limit the risk of starvation when the food supply is unpredictable (Anselme & Güntürkün, 2018). Previous studies of the influence of uncertainty on sign-tracking have focused on the uncertainty of whether or what outcomes will be presented by manipulating reward probability or magnitude. However, there can also be uncertainty about when an outcome will be available. One could introduce uncertainty about the timing of when a cue will be presented or uncertainty about when an outcome will be presented. One pigeon study manipulated the variability of cue durations and ITI durations at the same time (van Hest et al., 1986) and found that uncertainty about when a reward will be presented increased sign-tracking compared to when rewards were presented at a fixed time.

As mentioned above, it is also possible that uncertainty about when the cues for reward might appear could also affect sign-tracking. Here we manipulated the temporal uncertainty about when a cue will be presented by comparing fixed and variable ITIs. We found that, similar to other types of uncertainty, uncertainty in the timing of cue onset also increases sign-tracking. Additionally, increased sign-tracking has a negative effect on reward retrieval.We demonstrated that this increase is learned during initial exposure to the protocols and persists into extinction, even when temporal variability is reduced during the extinction phase, suggesting that the impact of temporal uncertainty on sign-tracking is learned rather than a transient effect on performance. These results suggest that the temporal predictability of cues strongly modulates cue-directed behaviors, and may play a critical role in shaping reward-related responses that are relevant to understanding addiction and compulsive behaviors.

## Method

### Animals and Housing Conditions

Male and female C57BL/6J wild-type mice (Mus musculus) were obtained from Jackson Laboratories (Bar Harbor, Maine). All animals were housed in a temperature- and humidity-controlled vivarium with a 12-h light/dark cycle and underwent behavioral testing in the light cycle. Mice were housed by sex in groups of 4 for the experiment. During behavioral testing, mice were subjected to food restriction to maintain between 85% and 90% of their ad lib diet body weight. Water was always available ad lib. Mice were at 14 weeks of age at the start of behavioral experiments. There were 32 subjects, 16 of each sex. All animal procedures were performed in accordance with the New York State Psychiatric Institute’s animal care and use committee’s regulations.

### Behavior Apparatus

Experimental chambers (22 x 18.1 x 12.7 cm) (Med-Associates, Vermont, Model: ENV-307W) had metal grid floors, two modular front and back walls, and two plexiglass walls. On one wall a centrally located food magazine (2.2 cm diameter) was equipped with a light and with an infrared head entry detector. A liquid dipper inside the food magazine delivered 0.01cc of evaporated milk. On either side of the food receptacle were retractable levers (1.59 x 0.79 cm) Both levers had a panel containing small green, yellow and red cue lights mounted above them (Med Associates Tripole LED panel model : ENV-322W). White house lights were mounted on the left side of the wall and were turned on during all sessions. Each chamber had an exhaust fan that generated white background noise at approximately 85 dBA. MedPC® software (Tatham & Zurn, 1989) was used to run the programs and collect the data with 10ms resolution.

### Acquisition Phase

We pre-trained the mice so that they were accustomed to the experimental setting and to obtaining rewards from a food magazine on a random interval schedule, with an average of 108 seconds between each reward for one session. To reduce contextual conditioning of this pre-training, we exposed the animals to background extinction on the following day without rewards to reduce the impact of magazine training on subsequent acquisition.

After pre-training and background extinction, we began the acquisition sessions. The mice were randomly assigned to one of four groups (n = 8 per group, 4 male and 4 female) based on the combination of two different durations of inter-trial interval (ITI) (72 s or144 s) and two levels of temporal uncertainty (fixed duration ITI or variable duration ITI). In the "fixed” groups, CSs were presented after a fixed duration from the end of a reward until the next CS was presented. In the "variable” groups, trials were presented after variable durations, with each ITI duration selected from a list of durations (4.36s to 227.11s for 72 s group, 8.72s to 454.21s for 144 s group) that were distributed according to an approximately exponential function such that the average duration matches that of the corresponding fixed group.

In each acquisition session, mice were exposed to a daily sequence of 24 trials. In all groups, each trial began with a 16-second presentation of the left lever (Conditioned Stimulus, CS), accompanied by the simultaneous illumination of green, yellow and red cue lights above the lever. Coincident with the termination of the 16 s lever presentation (lever retraction) and cue light, a dipper with 0.01 cc of evaporated milk (Unconditioned Stimulus: US) was presented for 5 seconds while the feeder light was activated to illuminate the food receptacle. These sessions took place five days a week for a total of 14 days. Throughout these sessions, both head entries into the trough and lever presses were automatically recorded.

### Extinction Phase

.After a 40-day break in training during which mice were given Ad Lib food and water, the same mice (n = 30; one mouse from the 72s-var group and one mouse from the 144s-fix group died before the start of the extinction phase) were retrained under the same condition they experienced during the acquisition phase until the sign-tracking and goal-tracking behaviors stabilized, totaling 6 sessions. For a subsequent extinction phase, each acquisition group was divided into two groups, half of which underwent extinction trials with temporal uncertainty (variable ITI), and the other half had temporal certainty (fixed ITI). To ensure that the new conditions were equally novel for all groups, an ITI length of 108 s (an average of the 144 s and 72 s ITI durations used during acquisition), was used in all extinction sessions, totaling 4 sessions. Half of each group was tested on extinction with a fixed 108 s ITI and the remainder were tested with a variable ITI that averaged 108 s. Thus, half of the subjects from each previous group were tested with a different ITI variability than the one they were originally trained with. This gave us the opportunity to see whether ITI variability affected initial learning and/or performance. As in acquisition, each lever extension lasted for 16 s accompanied by the simultaneous illumination of green, yellow and red cue lights above the lever, but no reward was presented after the CS.

### Data Analysis

Lever presses and head entries were recorded automatically by MedPC® hardware and organized using MATLAB. We used lever pressing during CS as an indicator of sign-tracking behavior, and head entries into the magazine (where the reward was dispensed) during CS as an indicator of goal-tracking as has been done in many previous studies (e.g. Boakes, 1977). The three measures of the PCA scores–sign-tracking score (or the response bias), the latency score, and the probability difference were calculated as previously described (Meyer et al., 2012); The sign-tracking score was calculated as (the number of LP - the number of HE) / (the number of LP + the number of HE), the latency score as: (HE latency during CS - LP latency during CS) / 16, and the probability difference as: (# of trials with at least one LP-the number of trials with at least one HE) / 24, where LP represents lever presses, and HE represents head entries into the magazine. All three scores range from -1.0 to +1.0, with -1.0 indicating all responses are goal-tracking (head entry) and +1.0 indicating all responses are sign-tracking (lever press). Positive scores means more sign-tracking behavior than goal-tracking behavior, and negative scores mean the opposite. The proportion of reward collected was calculated as (the number of trials where the animal’s head entered the magazine at least once during reward delivery) / (the total number of trials).

### Statistics

Statistical analyses were conducted using JASP Version 0.19.0 for Windows (Computer software, https://jasp-stats.org) Traditional statistical tests, including ANOVA, were employed to evaluate the effects of temporal uncertainty, ITI duration, and sex on the lever-pressing rate, head-entry rate, and PCA scores. Whenever sphericity assumptions were violated, the Geisser-Greenhouse correction was applied prior to conducting repeated measures ANOVA to ensure the validity of the results. Post-hoc tests were conducted using Bonferroni correction when appropriate. Details of the statistical analyses are provided as tables.

### Transparency and Openness

The datasets generated and the custom code used for analysis in the current study are available at an online repository at https://osf.io/y4v62/?view_only=48497c0dd5b44012a0f470f572b8704b. This experiment was not pre-registered.

## Results

### Temporal Uncertainty About Cue Onset Increases Sign Tracking During Acquisition

Previous studies have documented the impact of ITI length on sign-tracking, with longer ITIs shown to increase sign-tracking behavior (Gibbon et al., 1980; Lee et al., 2018; Mahmoudi et al., 2023). To further explore the effect of ITI length and its interaction with ITI variability, we investigated differences between groups across four different ITI conditions characterized by combinations of ITI duration (72 seconds, 144 seconds) and ITI variability (fixed, varied). Sign-tracking behavior increased over blocks (2 sessions per block) (Figure. 1A), while goal-tracking generally decreased (Figure. 1B), resulting in a significant rise in the sign-tracking scores. This direction of effects was observed across all groups. The groups with the variable ITIs exhibited more sign-tracking behavior than the fixed ITI groups. After applying Geisser-Greenhouse correction, a three-way ANOVA (Block * ITI variability * ITI length) was conducted on sign-tracking behavior (lever press), goal-tracking behavior (head entry), and PCA scores. For lever pressing, There were significant main effects of Block (p < 0.001), ITI variability (p = 0.009), but no main effect of ITI length (p = 0.059). Post-hoc tests revealed significant increases in lever pressing across blocks, with this development observed early in training (Block 1 vs Block2, Block 3, Block4: 0.044, p<0.004, p<0.001, respectively). Head entries were significantly affected only by blocks (p = 0.002), and according to post-hoc tests, the development of head entries progressed more slowly than lever pressing (Block 1 vs Block 7, p = 0.0314).

**Figure 1.**
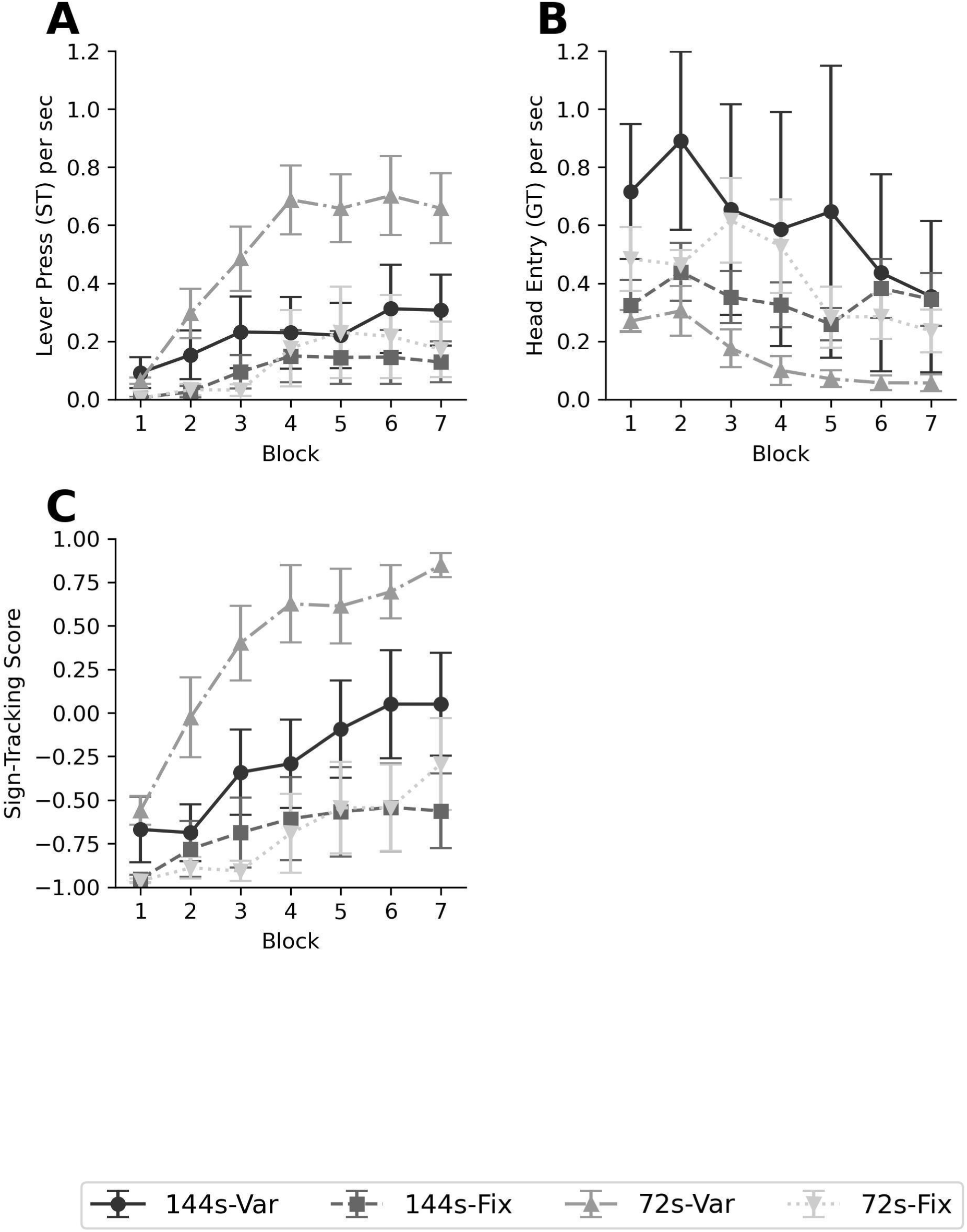
Conditioned Responses during acquisition phase by group (144s-Var, 144s-Fix, 72s-Var, 72s-Fix; n = 8 per group). *Note.* Response rates were measured during the CS (16s) that was followed by reward. *(A)* Sign-tracking rate during acquisition: the number of lever presses per second. (B) Goal-tracking rate during acquisition: the number of head entries into the food magazine per second. (C) Sign-tracking Score: defined as the ratio of bias toward ST behavior relative to the total response. Calculated as: [(Number of lever presses - Number of head entries) / (Number of lever presses + Number of head entries)]. Each block consists of the average value from two consecutive sessions. Datapoints represent the group mean, Error bars represent the standard error (SE) of the mean.

All three components of the PCA score—sign-tracking score, latency score, and probability difference—exhibited significant main effects of Block (p<0.001 for Sign-tracking score, latency score, and probability difference) and ITI variability (p < 0.001 for sign-tracking score and probability difference, p =0.003 for latency score) (Figure. 1C, Supplemental Figure. 1A, B). Post-hoc tests with Bonferroni correction revealed that the effect of Block consistently emerged at the same point in training—Block 3 compared to Block 1—for all measures (p = 0.002, p = 0.0021, p = 0.009, respectively). Only probability difference exhibited a Block × Variability interaction (p=0.049), and post-hoc analysis showed that this interaction was driven by a more pronounced score increase in the variable group between Blocks 2 and 3 (p=0.043). Across acquisition, sex had no significant effect on any of the measures of conditioned responding (Supplemental Table 1). In summary, training induced the development of sign-tracking behaviors in all groups, with sign-tracking developing more rapidly than goal-tracking. Proportion of rewards collected was not affected by either ITI variability or ITI length (Supplemental Table 2).

### Extinction of Conditioned Responses

After a break in the experiment, mice were retrained for six days under the same protocol they experienced during the acquisition phase. After 6 days of retraining, sign-tracking and goal-tracking behaviors were stable across days and there was no difference between the sign-tracking and goal-tracking measures from the last block of the acquisition phase and the average of the 6 retraining sessions, Paired t-tests: (Head entry: [t(29) = 0.570, p = 0.573]; lever press: [t(29) =1.010, p = 0.321]) (Data not shown).

During the extinction phase, all mice experienced a fixed ITI of 108 seconds (intermediate between the 72s and 144s ITIs used during the acquisition phase) over the course of four extinction sessions. Half the mice in each of the 4 acquisition groups experienced variable ITIs and half experienced fixed ITIs. When analyzing the mice by extinction groups only (without regard for acquisition conditions), there was no main effect of ITI variability on goal-tracking, sign-tracking, or PCA scores (Figure 2, Supplemental Figure 2A, 2B). Due to differing levels of conditioned responses prior to the extinction phase, we conducted a two-way repeated measures ANCOVA (Block*Extinction group, covariate = Pre-Extinction values). The analysis revealed no main effect of the extinction session on either sign-tracking (p = 0.179) or goal-tracking (p = 0.763), indicating that changes across all extinction sessions on average were not robust. However, pairwise comparisons demonstrated significant effects in the earlier stages of the extinction phase. Specifically, significant differences were observed between Session 1 and Session 2 for both sign-tracking and goal-tracking ([t =8.186, p < 0.001], [t = 4.633, p <0.01], respectively), Session 1 and Session 3 ([t = 4.494, p < 0.001], [t = 10.321, p <0.01], respectively), and Session 1 and Session 4 ([t = 3.633, p < 0.008], [t = 10.243, p < 0.001], respectively), but no significant changes are detected between session 3 and session 4 for either sign-tracking and goal-tracking. These results indicate a clear decline in conditioned responses early in the extinction phase, with little to no further decline observed later in the phase. There were also no effects of the extinction group and no significant interaction between session and extinction group for either sign-tracking or goal-tracking. This suggests that ITI variability during extinction had no effect on the extinction of conditioned responses.

**Figure 2.**
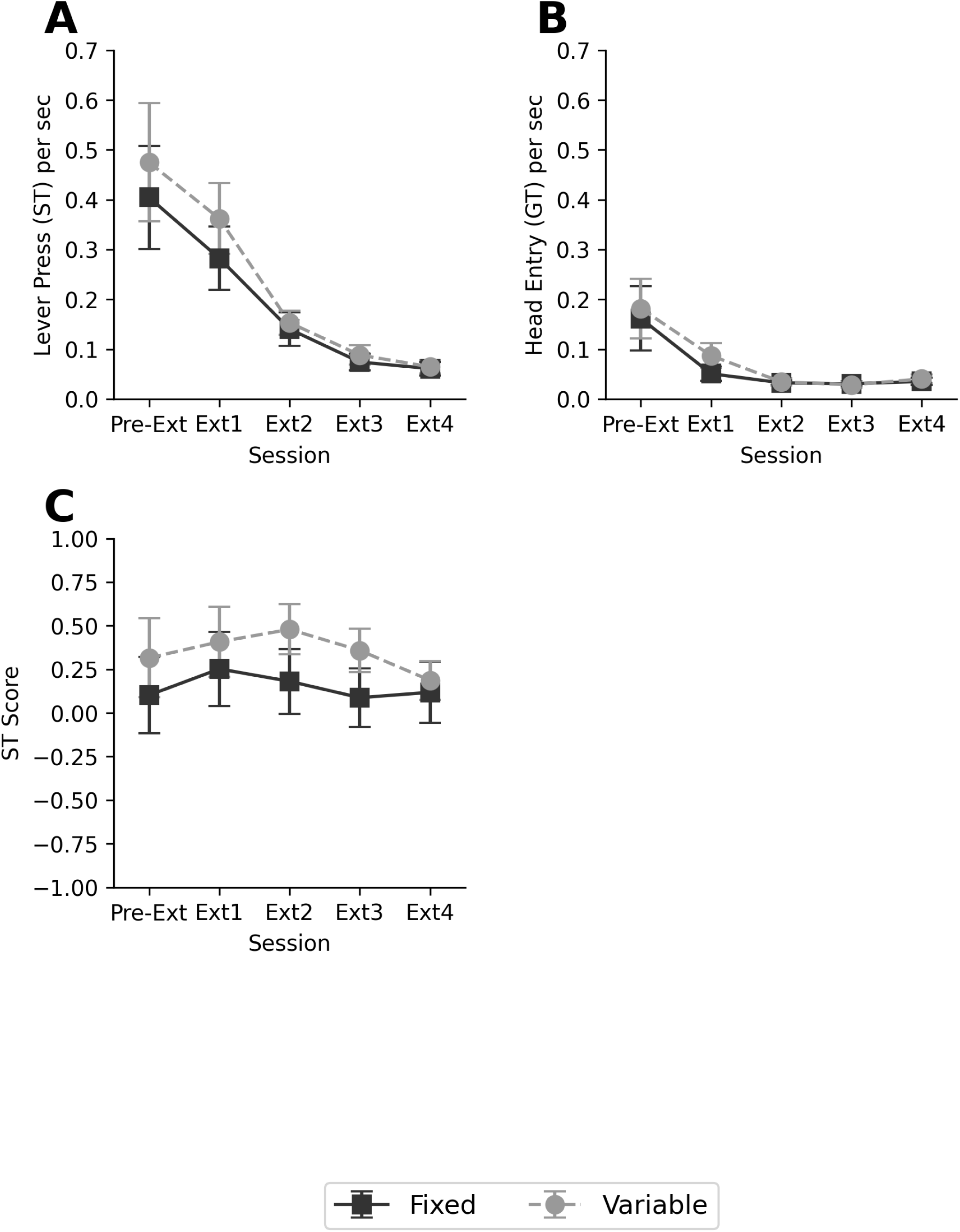
Conditioned Responses Across Extinction Sessions by Extinction Groups (108s-Fix and 108s-Var; n = 15 per group) *Note.* Response rates are measured during the CS (16 s), which was not followed by reward. The leftmost point on the x-axis represents the average of the last two sessions of retraining before the start of the extinction phase. (A) Goal-tracking Rate: Number of head entries per second. (B) Sign-tracking Rate: Number of lever presses per second. (C) Sign-tracking Score: Defined as the ratio of bias toward ST behavior relative to the total response. The ST score is calculated as: (Number of lever presses - Number of head entries) / (Number of lever presses + Number of head entries). Data points represent group mean and error bars indicate the standard error (SE) of the mean.

To assess whether the tendency to engage in sign-tracking or goal-tracking shifted during extinction, we conducted a two-way ANCOVA on the sign-tracking scores, latency scores, and probability differences. No significant effect of the extinction group was observed for any of these measures. Session effects were also not significant for the sign-tracking score or latency score but were significant for the probability difference (p < 0.001) with the significant decline later in the sessions (Session 2 and Session 3 [t = 3.870, p = 0.004]; session 3 and session 4 [t = 3.211, p = 0.020]). These results suggest that overall, sign-tracking and goal-tracking declined at similar rates during extinction. In summary, unlike during the acquisition phase, ITI variability did not influence the rate of extinction of animals’ conditioned responses. Both head entries and lever presses were extinguished at comparable rates.

Analyzing the extinction data according to conditions experienced during the acquisition phase, irrespective of extinction condition, revealed that the groups that experienced the variable ITI during acquisition continued to do more sign tracking and less goal tracking than groups that experienced the fixed ITI during acquisition. Three-way repeated measures ANCOVAs (ITI length during acquisition * ITI variability during acquisition * extinction session; Pre-Extinction levels as covariate) were conducted. A main effect of ITI variability during acquisition was observed in sign-tracking (p < 0.001), sign-tracking score (p = 0.045), latency score (p = 0.029), and probability difference (p = 0.003), confirming the consistent effect of variability during the acquisition phase on the extinction process (Figure 3A, Supplemental Figure 3A, 3B). The interaction of variability during acquisition and session was observed only in probability score (p = 0.011). This interaction indicates a consistent decline in the probability score in the variable ITI group compared to the relatively stable probability score in the fixed ITI group, likely due to the higher initial probability score in the variable group. ITI variability during acquisition did not affect goal-tracking. Altogether, these results indicate a robust and persistent effect of ITI variability during acquisition on the level of sign-tracking behavior and sign-tracking tendencies throughout extinction. This effect occurred independently of ITI variability during the extinction phase itself (Figure 3).

**Figure 3.**
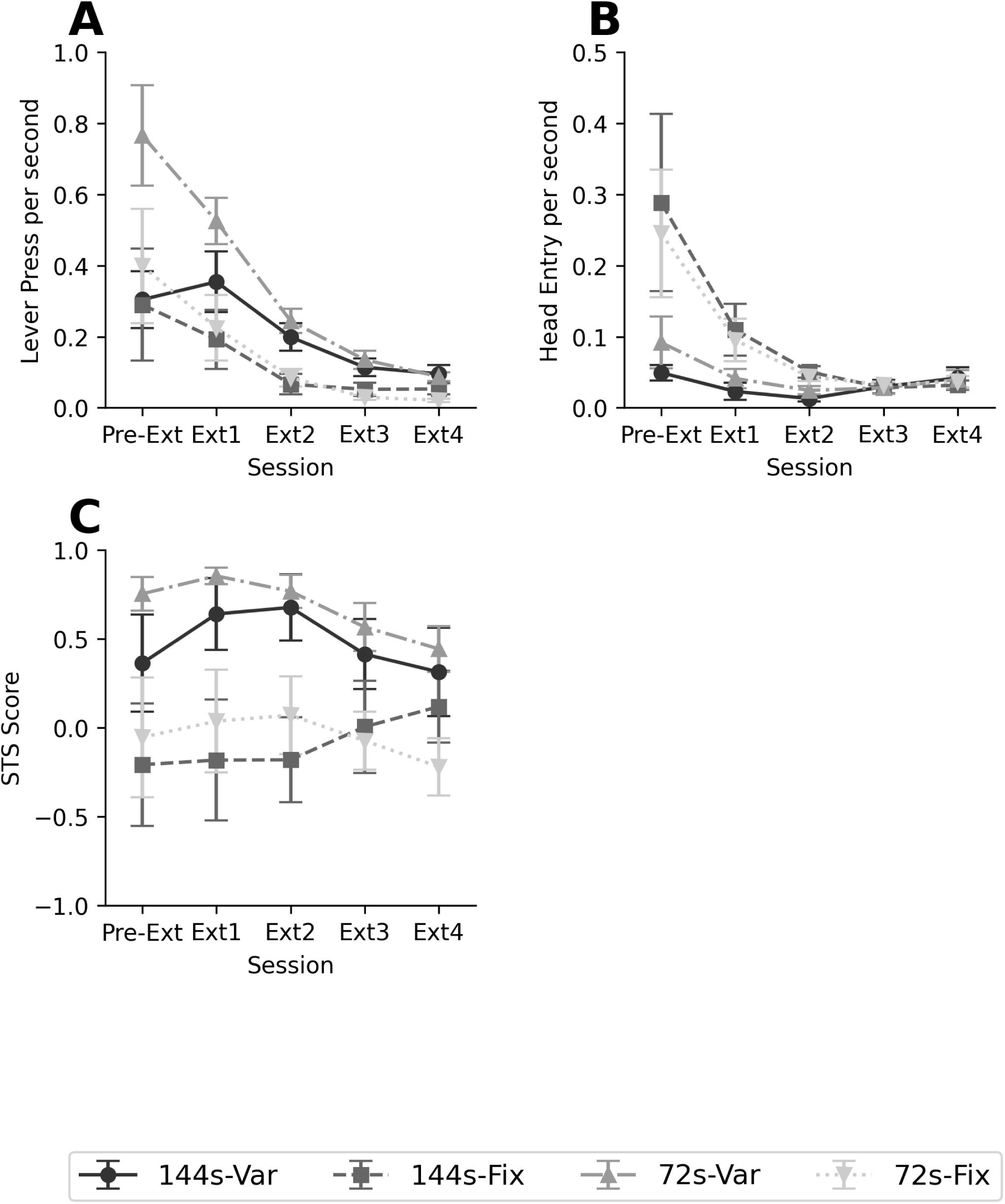
Conditioned responses over extinction sessions by acquisition groups. (144s-Var: n = 8, 144s-Fix: n = 7, 72s-Var: n = 8, 72s-Fix: n = 8) *Note.* The leftmost point on the x-axis is the average of the last two sessions of retraining before the start of the extinction phase. (A) Sign-tracking rate (The number of lever presses per second). (B) Goal-tracking rate (The number of head entries per second) (C) Sign-tracking Score: Defined as the ratio of bias toward ST behavior relative to the total response. The ST score is calculated as: (Number of lever presses - Number of head entries) / (Number of lever presses + Number of head entries). Data points represent group mean and error bars indicate the standard error (SE) of the mean.

## Discussion

Here we observed a general increase in sign-tracking behavior across acquisition sessions in all groups, with a more pronounced increase among those exposed to variable inter-trial intervals (ITI) during acquisition. This suggests that variable ITIs enhance the incentive salience of cues associated with rewards. Interestingly, while ITI variability during the *<u>extinction phase</u>* did not directly influence the rate at which conditioned responses were extinguished, ITI variability during the *<u>acquisition phase</u>* continued to have a significant impact. Mice that experienced variable ITIs during acquisition maintained higher levels of sign-tracking across all measurements during the extinction sessions. This finding indicates a stronger and more persistent bias toward the cue over the reward, even as extinction progressed. This effect underscores that early exposure to temporal uncertainty can have lasting effects on subsequent extinction of sign-tracking. This demonstrates that the impact of ITI variability observed in the acquisition phase was a learned effect, not merely a performance effect based on the current conditions the animals were experiencing.

In this study, the effect of ITI length on sign-tracking behavior or sign-tracking scores between subjects did not reach statistical significance. This finding contrasts with previous studies (Gibbon et al., 1980; Lee et al., 2018; Mahmoudi et al., 2023; Terrace et al., 1975), which reported that longer ITIs increased sign-tracking. There are many differences between the current experiment and previous studies including the ITI and CS durations as well as the specific distributions of ITI durations. Additionally, the subjects of the current experiment were mice but in previous studies pigeons (Gibbon et al., 1980; Terrace et al., 1975) or rats (Lee et al., 2018; Mahmoudi et al., 2023) were studied. Any or all of these differences may have contributed to the difference between our study and previous ones with respect to the effect of ITI duration.

Our finding that temporal uncertainty due to ITI variability enhances sign-tracking behavior aligns with numerous previous studies showing that other forms of uncertainty similarly enhance sign-tracking (Davey & Cleland, 1982; Anselme et al., 2013). Previous research has primarily explored uncertainty regarding reward properties, such as whether a reward is delivered or not, or reward magnitude (Anselme et al., 2013; Monosov & Hikosaka, 2013). In contrast, our study specifically focused on uncertainty about *when* a *cue* will be presented. Similar to other forms of uncertainty, temporal uncertainty regarding the cue had a comparable effect on sign-tracking behavior. This suggests that the critical factor for sign-tracking is the uncertainty surrounding the CS-US pairing rather than the uncertainty in the characteristics of the reward signaled by the cue itself. In other words, even when the reward signaled by the CS is known, if the timing of the CS or the CS-US pairing is uncertain, incentive salience is attributed to the cue, even when it has full predictive value.

These results demonstrate that temporal uncertainty in cue presentation leads to greater sign-tracking. The lasting impact of this uncertainty during extinction suggests that the modulation of cue-directed behaviors by temporal factors is a robust and learned effect. The enduring influence of initial conditioning,, may contribute to the difficulty of extinguishing cues that trigger addictions and compulsions in real-world contexts.

**Table 1.**
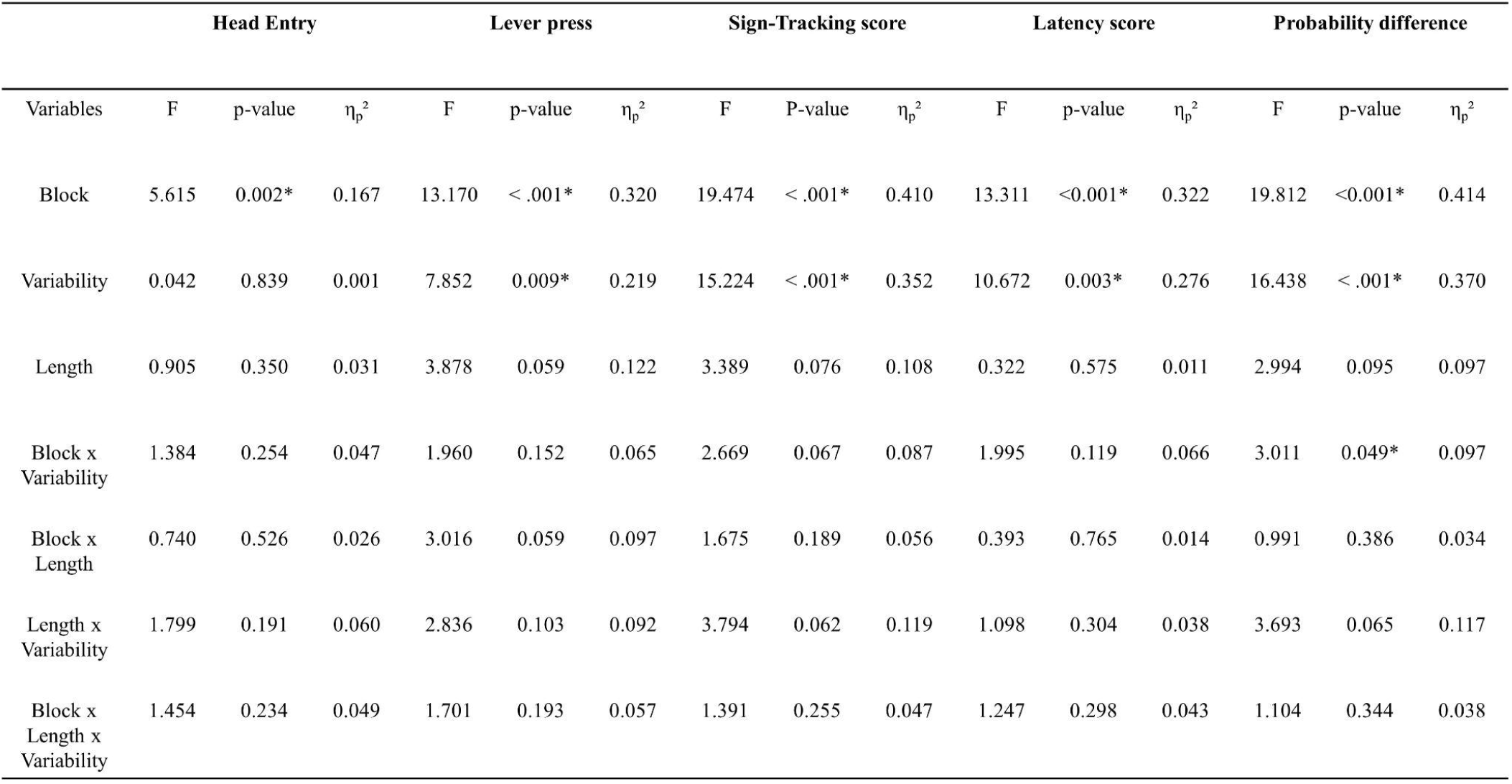
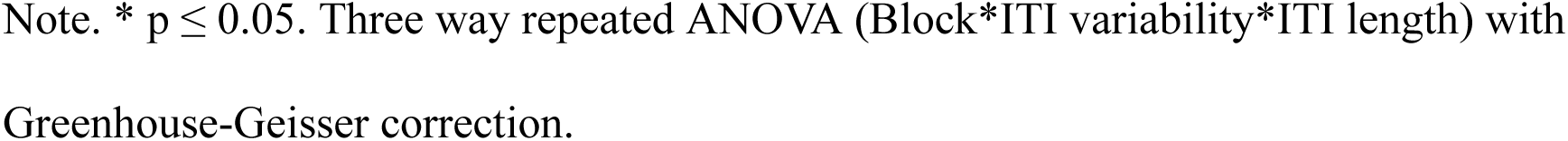
Summary statistics for acquisition phase.

*Summary statistics for acquisition phase*
| Variables | Head Entry |  |  | Lever press |  |  | Sign-Tracking score |  |  | Latency score |  |  | Probability difference |  |  |
| --- | --- | --- | --- | --- | --- | --- | --- | --- | --- | --- | --- | --- | --- | --- | --- |
| | F | p-value | $\eta_p^2$ | F | p-value | $\eta_p^2$ | F | P-value | $\eta_p^2$ | F | p-value | $\eta_p^2$ | F | p-value | $\eta_p^2$ |
| Block | 5.615 | 0.002* | 0.167 | 13.170 | < .001* | 0.320 | 19.474 | < .001* | 0.410 | 13.311 | <0.001* | 0.322 | 19.812 | <0.001* | 0.414 |
| Variability | 0.042 | 0.839 | 0.001 | 7.852 | 0.009* | 0.219 | 15.224 | < .001* | 0.352 | 10.672 | 0.003* | 0.276 | 16.438 | < .001* | 0.370 |
| Length | 0.905 | 0.350 | 0.031 | 3.878 | 0.059 | 0.122 | 3.389 | 0.076 | 0.108 | 0.322 | 0.575 | 0.011 | 2.994 | 0.095 | 0.097 |
| Block x Variability | 1.384 | 0.254 | 0.047 | 1.960 | 0.152 | 0.065 | 2.669 | 0.067 | 0.087 | 1.995 | 0.119 | 0.066 | 3.011 | 0.049* | 0.097 |
| Block x Length | 0.740 | 0.526 | 0.026 | 3.016 | 0.059 | 0.097 | 1.675 | 0.189 | 0.056 | 0.393 | 0.765 | 0.014 | 0.991 | 0.386 | 0.034 |
| Length x Variability | 1.799 | 0.191 | 0.060 | 2.836 | 0.103 | 0.092 | 3.794 | 0.062 | 0.119 | 1.098 | 0.304 | 0.038 | 3.693 | 0.065 | 0.117 |
| Block x Length x Variability | 1.454 | 0.234 | 0.049 | 1.701 | 0.193 | 0.057 | 1.391 | 0.255 | 0.047 | 1.247 | 0.298 | 0.043 | 1.104 | 0.344 | 0.038 |

**Table 2.** Summary statistics for extinction phase (extinction session*ITI variability during extinction)

|  | Head Entry |  |  | Lever press |  |  | Sign-Tracking score |  |  | Latency score |  |  | Probability difference |  |  |
| --- | --- | --- | --- | --- | --- | --- | --- | --- | --- | --- | --- | --- | --- | --- | --- |
| Variables | F | p-value | $\eta_p^2$ | F | p-value | $\eta_p^2$ | F | P-value | $\eta_p^2$ | F | p-value | $\eta_p^2$ | F | p-value | $\eta_p^2$ |
| Session | 0.725 | 0.474 | 0.026 | 1.824 | 0.179 | 0.063 | 1.602 | 0.205 | 0.056 | 1.329 | 0.272 | 0.047 | 11.624 | < 0.001* | 0.301 |
| Ext-variability | 0.874 | 0.358 | 0.031 | 0.257 | 0.617 | 0.009 | 0.378 | 0.544 | 0.014 | 0.107 | 0.746 | 0.004 | 0.254 | 0.618 | 0.009 |
| Ext-variability x Session | 2.581 | 0.092 | 0.087 | 0.734 | 0.458 | 0.026 | 1.339 | 0.270 | 0.047 | 1.283 | 0.287 | 0.045 | 0.331 | 0.756 | 0.012 |

Note. \* $p \leq 0.05$ . Two way repeated ANOVA (Session\*ITI variability during extinction; Pre-Extinction values as covariate) with Greenhouse-Geisser correction. Pre-Extinction values are the average of the last two sessions of the retraining phase.

**Table 3.** Summary statistics for extinction phase (extinction session*ITI variability during acquisition * ITI length during acquisition)

IMPACT OF TEMPORAL UNCERTAINTY ON SIGN-TRACKING BEHAVIOR
| Variables | Head Entry |  |  | Lever press |  |  | Sign-Tracking score |  |  | Latency score |  |  | Probability difference |  |  |
| --- | --- | --- | --- | --- | --- | --- | --- | --- | --- | --- | --- | --- | --- | --- | --- |
| | F | p-value | $\eta_p^2$ | F | p-value | $\eta_p^2$ | F | P-value | $\eta_p^2$ | F | p-value | $\eta_p^2$ | F | p-value | $\eta_p^2$ |
| Acquisition Variability | 0.509 | 0.482 | 0.020* | 16.591 | < .001* | 0.399 | 4.460 | 0.045* | 0.151 | 5.385 | 0.029* | 0.177 | 11.180 | 0.003* | 0.309 |
| Acquisition Length | 0.102 | 0.752 | 0.004* | 0.458 | 0.505 | 0.018 | 0.460 | 0.504 | 0.018 | 0.011 | 0.919 | 0.0004 | 0.131 | 0.721 | 0.005 |
| Session x Acquisition Variability | 3.327 | 0.054 | 0.117 | 2.417 | 0.113 | 0.088 | 1.361 | 0.264 | 0.052 | 1.797 | 0.163 | 0.067 | 4.507 | 0.011* | 0.153 |
| Session x Acquisition Length | 0.003 | 0.992 | 0.0001* | 0.439 | 0.600 | 0.017 | 1.476 | 0.233 | 0.056 | 0.815 | 0.477 | 0.032 | 0.094 | 0.937 | 0.004 |

Note. \* $p \leq 0.05$ . Three way repeated ANCOVA (Session\*ITI variability during acquisition\* ;
Pre-Extinction values as covariate) with Greenhouse-Geisser correction. Pre-Extinction values are the average of the last two sessions of the retraining phase.

---

**Supplemental Table 1.** Statistics including sex as variable.

| Variables | Head Entry |  |  | Lever press |  |  | Sign-Tracking score |  |  | Latency score |  |  | Probability difference |  |  |
| --- | --- | --- | --- | --- | --- | --- | --- | --- | --- | --- | --- | --- | --- | --- | --- |
| | F | p-value | $\eta_p^2$ | F | p-value | $\eta_p^2$ | F | P-value | $\eta_p^2$ | F | p-value | $\eta_p^2$ | F | p-value | $\eta_p^2$ |
| Sex | 1.147 | 0.295 | 0.046 | 0.006 | 0.937 | 0.0003 | 0.107 | 0.746 | 0.004 | 0.102 | 0.752 | 0.004 | 0.003 | 0.958 | 0.0001 |
| Sex x Block | 0.381 | 0.733 | 0.016 | 0.187 | 0.824 | 0.008 | 0.189 | 0.855 | 0.008 | 1.801 | 0.151 | 0.070 | 0.080 | 0.934 | 0.003 |
| Sex x Variability | 1.508 | 0.231 | 0.059 | 0.103 | 0.751 | 0.004 | 0.292 | 0.594 | 0.012 | $7.306 \times 10^{-5}$ | 0.993 | $3.04 \times 10^{-6}$ | 0.090 | 0.766 | 0.004 |
| Sex x Length | 2.218 | 0.149 | 0.085 | 0.670 | 0.421 | 0.027 | 0.316 | 0.579 | 0.013 | 1.241 | 0.276 | 0.049 | 0.419 | 0.523 | 0.017 |
| Sex x Variability x Length | 0.547 | 0.467 | 0.022 | 0.0005 | 0.982 | $2.2 \times 10^{-5}$ | 0.046 | 0.832 | 0.002 | $7.261 \times 10^{-5}$ | 0.979 | $3.03 \times 10^{-5}$ | 0.0005 | 0.982 | $2.20 \times 10^{-5}$ |
| Sex x Variability x Length x Block | 1.165 | 0.326 | 0.046 | 0.083 | 0.916 | 0.003 | 0.552 | 0.602 | 0.022 | 0.253 | 0.869 | 0.010 | 1.270 | 0.291 | 0.050 |

**Supplemental Table 2.** Statistics of proportion of reward collected.

IMPACT OF TEMPORAL UNCERTAINTY ON SIGN-TRACKING BEHAVIOR
| Proportion of reward collected |  |  |  |
| --- | --- | --- | --- |
| Variables | F | p-value | $\eta_p^2$ |
| Block | 2.714 | 0.049* | 0.088 |
| Variability | 0.003 | 0.959 | 9.705 x 10<sup>-5</sup> |
| Length | 0.358 | 0.555 | 0.013 |
| Block x Variability | 0.595 | 0.623 | 0.021 |
| Block x Length | 0.915 | 0.439 | 0.032 |
| Block x Variability x Length | 1.306 | 0.278 | 0.045 |

**Supplemental Figure 1.**
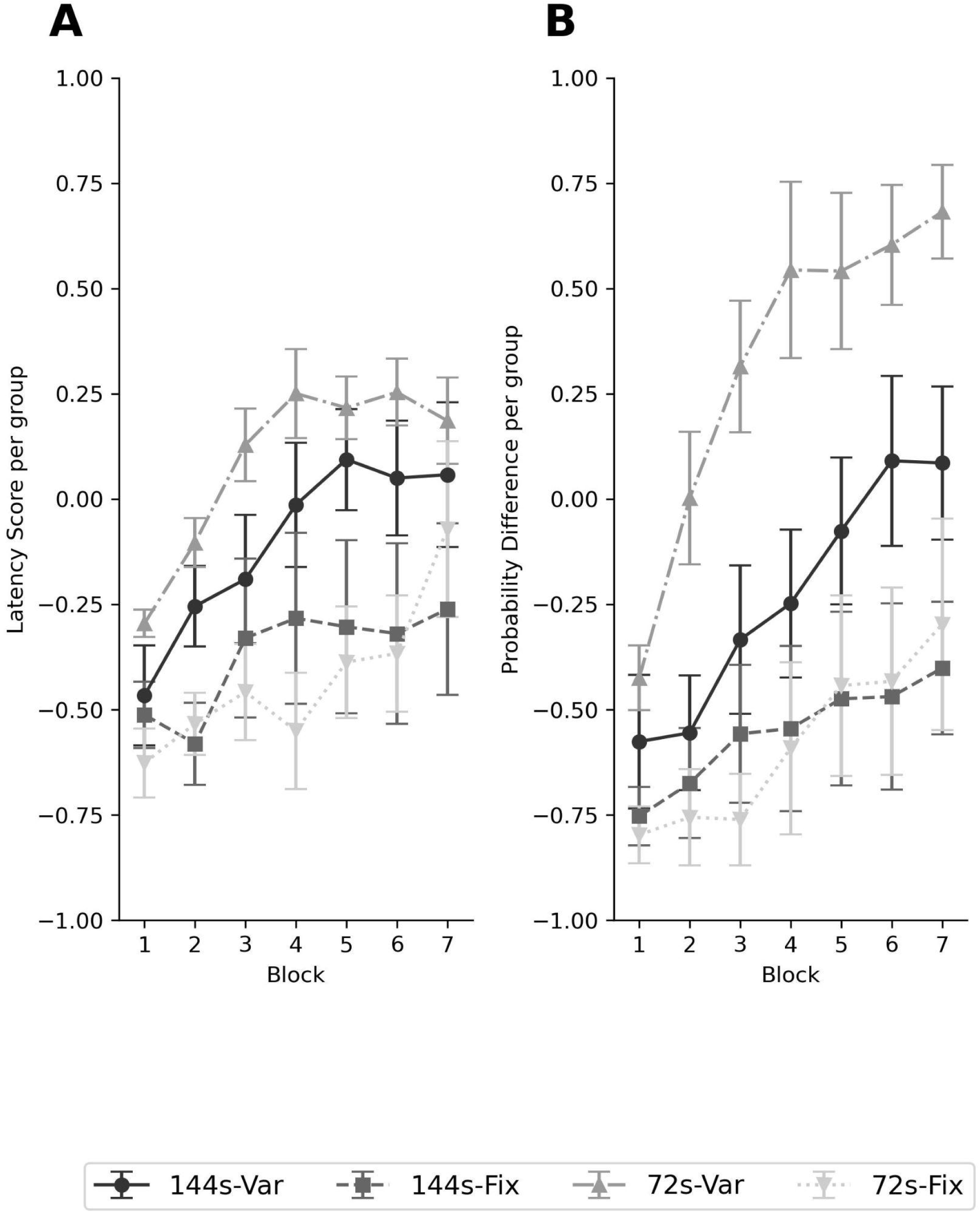
Latency score and probability difference during acquisition phase by group (144s-Var, 144s-Fix, 72s-Var, 72s-Fix; n = 8 per group). *Note.* (A) Latency score during acquisition. Calculated as: [(Head Entry latency during CS -Lever Press Latency during CS) / (The number of trials per session)]. Each block consists of the average value from two consecutive sessions. Datapoints represent the group mean, Error bars represent the standard error (SE) of the mean.

**Supplemental Figure 2.**
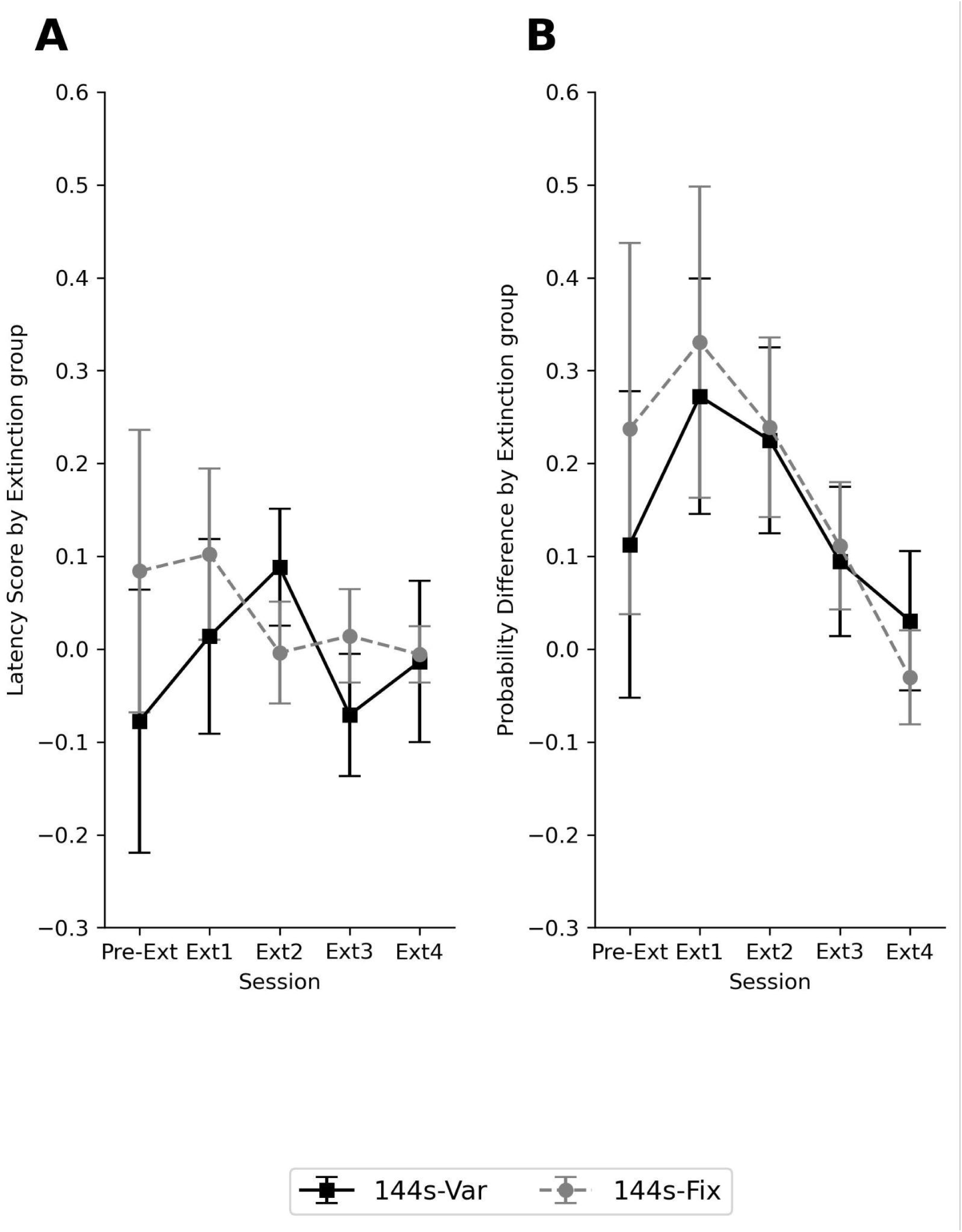
Latency score and probability difference during extinction phase by extinction group (108s-Var, 108s-Fix; n = 15 per group). *Note.* (A) Latency score during acquisition. calculated as: [(Head Entry latency during CS -Lever Press Latency during CS) / (The number of trials per session)]. Each block consists of the average value from two consecutive sessions. Datapoints represent the group mean, Error bars represent the standard error (SE) of the mean.

**Supplemental Figure 3.**
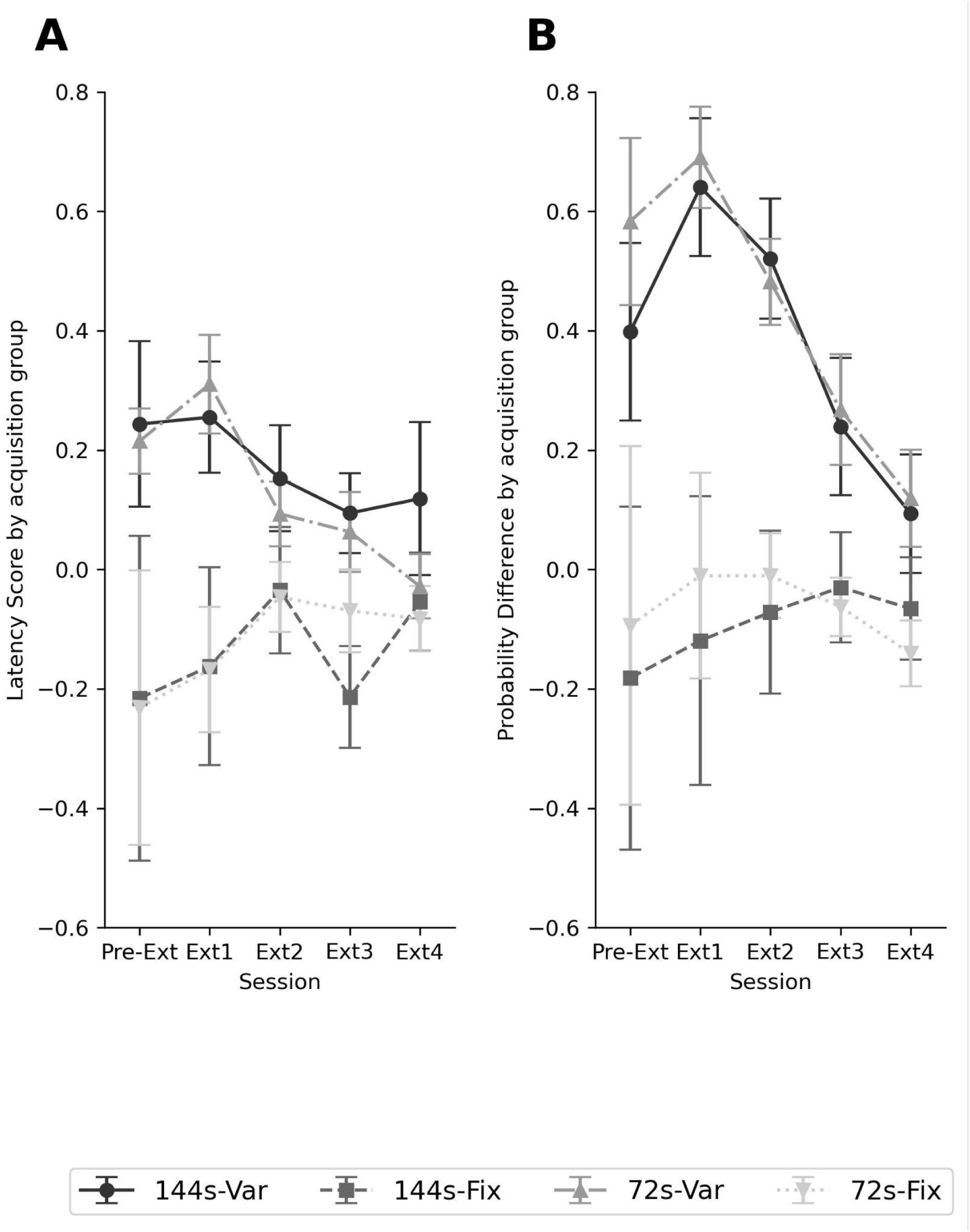
Latency score and probability difference during extinction phase by acquisition group (144s-Var, 144s-Fix, 72s-Var, 72s-Fix). *Note.* (A) Latency score during acquisition. calculated as: [(Head Entry latency during CS -Lever Press Latency during CS) / (The number of trials per session)]. Each block consists of the average value from two consecutive sessions. Datapoints represent the group mean, Error bars represent the standard error (SE) of the mean.

